# Melanoma suppresses galectin-9-glycan axis in dendritic cells and galectin-9 restoration limits T regulatory cell expansion

**DOI:** 10.64898/2026.09.09.749950

**Authors:** Andrea Rodgers-Furones, Amaia González De Zárate, Theodoros Basiakos, Colin Lee, Guido van Mierlo, Karina Valeria Mariño, Laia Querol Cano

## Abstract

Dendritic cells (DCs) are key orchestrators of anti-tumor adaptive immune responses. Their function is tightly regulated by galectins, a family of carbohydrate-binding proteins that decode extracellular glycans into intracellular signaling. Here, we show that exposure to melanoma-conditioned media (CM) induces loss of galectin-9 (gal-9) at the DC surface. Notably, gal-9 loss, independent of transcriptional regulation, was associated with the acquisition of an immunosuppressive phenotype in CD14⁺cDC2 cells following tumor exposure, with higher-molecular weight fractions of the melanoma secretome as mediators. Notably, gal-9 depletion was mirrored with a reduction in gal-9 ligands on the cell surface, particularly GalNAc-containing glycoepitopes, pointing towards a melanoma-exploited gal-9-glycan axis as a DC-evasive strategy. Interestingly, restoring gal-9 surface levels in CM-exposed CD14⁺cDC2 cells prevented the expansion of regulatory T cells (Tregs), postulating gal-9 as a novel immunomodulatory molecule in DC-mediated Treg induction during melanoma progression. Altogether, our data suggest that melanoma-derived factors remodel the DC glycan-gal-9 axis to enhance T cell differentiation towards regulatory phenotypes and dampen anti-tumor immunity. This identifies the gal-9/glycan axis in CD14⁺cDC2 cells as a vulnerable node in melanoma immune evasion and as a potential therapeutic target.

## Introduction

Dendritic cells (DCs) are essential for launching and shaping anti-melanoma immune responses by virtue of their ability to present tumor antigens to effector T cells^1^. They subdivide into conventional type 1 and 2 DCs (cDC1 and cDC2) and plasmacytoid DCs (pDCs). In human blood, CD1c^+^ cells are the most abundant subset. They excel at activating CD4^+^ T cells and skewing adaptive responses towards T-helper type 1 (Th)1/Th17-mediated immunity, crucial for protective immunity against intracellular pathogens and anti-tumor inflammatory responses^2^. In early-phase clinical trials, cDC2 vaccination in patients with solid tumors has been associated with enhanced cytotoxic T cell responses and signals of improved progression-free survival^3, 4, 5^. Importantly, cDC2s present high phenotypic and functional plasticity in response to soluble tumor microenvironment (TME)-related cues, such as IL-10, TGF-β, IL-6, CSF-1R, PGE₂, VEGF and hypoxia^6, 7, 8, 9, 10, 11^. This plasticity shapes both the generation of antitumor immune responses and the clinical efficacy of immunotherapies in melanoma patients^6, 7, 12, 13^. This is related to the acquisition of immunosuppressive features, most notably the upregulation of monocytic-lineage markers such as CD14 and CD163, and a shift in T cell priming towards T regulatory (Tregs)/Th2 rather than Th1/Th17, ultimately promoting dysfunctional, tolerized T cell responses^6, 14^.

Emerging evidence indicates that TME-inflammatory milieu can remodel the glycome of tumor-exposed immune cells toward an immunosuppressive phenotype, as shown for CD8⁺ T cells where altered N-glycosylation and increased MGAT5-dependent branching contributed to exhaustion, and for macrophages where changes in sialylation influenced tumor-associated macrophage polarization and response to checkpoint blockade^15, 16, 17^. Accordingly, glycan-related strategies are gaining interest in tumor immunology, with the sialoglycan-siglec axis emerging as a particularly prominent target^18, 19^. Galectins are soluble lectins that function as molecular “readers” of surface glycans on immune cells, translating information encoded by glycans into cellular signaling, ultimately fine-tuning cell function^20^. Moreover, galectins have been described as key modulators of immune responses, shaping leukocyte activation, differentiation, migration, and survival^20, 21, 22^. Despite their well-established immunomodulatory functions, whether and how tumors exploit galectin-glycan interactions to evade immune surveillance remains underexplored.

Galectin-9 (gal-9) is a tandem-repeat galectin composed of two carbohydrate-recognition domains (CRDs) joined by a flexible linker. At the cell surface, gal-9 recognizes and crosslinks β-galactoside-containing glycoconjugates, particularly poly-*N-*acetyl-Lactosamine (polyLacNAc) chains and A or B blood group antigens^23, 24^. Within the TME, gal-9 engages glycosylated immune checkpoint receptors such as Tim-3 and PD-1 on exhausted T cells and promotes their apoptosis, while also enhancing the suppressive activity of regulatory T cells, thereby contributing to defective antitumor immunity^25^. This has positioned gal-9 as a key regulator of anti-tumoral responses; consequently, therapeutic strategies to inhibit gal-9-mediated pathways are now being explored. In contrast and highlighting its pronounced cell-dependent pleiotropy, we and others have shown that gal-9 exerts immunostimulatory effects on DCs by supporting antigen phagocytosis, cytokine secretion, migration, maturation, and immunological synapse formation with T cells underscoring its relevance^26, 27, 28, 29^.

Gal-9 regulation in tolerogenic DCs remains poorly defined. We identifed a tumor-driven loss of gal-9 at the surface of immunosuppressive CD14⁺cDC2 populations. Mechanistically, melanoma-secreted high-molecular weight factors promote a transient, post-transcriptional reduction in surface gal-9 alongside changes in DC glycan features consistent with altered gal-9 binding. In turn, restoring gal-9 reprograms DC-T cell outcomes by limiting Treg expansion and enhancing IFN-γ⁺ effector responses, revealing a glycan-dependent mechanism by which melanoma constrains DC function.

## Materials and methods

### Generation of melanoma-conditioned medium

Conditioned medium (CM) was obtained from A375 cells, cultured in DMEM (Thermo Fisher Scientific, #10566016) supplemented with 10% fetal bovine serum (FBS) and antibiotic-antimycotic (Thermo Fisher Scientific, #15240096) and seeded at an initial density of 0.1x10^6^ cells/ml for 72 hours (37 °C, 5% CO₂). Cell medium was collected, spun down to remove dead cells and debris, aliquoted and stored at -20 °C until used.

Molecular weight fractionation of CM was performed using Amicon Ultra centrifugal filter devices (Merck, Darmstadt, Germany) according to manufacturer’s instructions. CM was sequentially filtered on devices with molecular weight cutoff of 100 kDa (#UFC910024), 50 kDa (#UFC805024) and 10 kDa (#UFC801024). To correct for the concentration of the CM components inherent to the fractionation protocol, CM fractions were resuspended with the appropriate volume of DMEM supplemented with 10% FBS and antibiotic-antimycotic to reach the initial volume prior to fractionation. Media were aliquots and stored at -20 °C until used.

### Isolation of primary cells

Primary cells were derived from peripheral blood mononuclear cells (PBMCs) of healthy blood donors with their informed consent (Sanquin, Nijmegen, the Netherlands). PBMCs were obtained via density gradient centrifugation with Lymphoprep density gradient medium (STEMCELL technologies, #18061) according to manufacturer’s instructions.

cDC2s were isolated using the magnetic activated cell sorting (MACS) CD1c^+^ isolation kit (Miltenyi, #130-119-475) according to manufacturer’s instructions. To isolate all DC subsets (panDCs), PBMCs were incubated with anti-CD14 (Miltenyi, #130-050-201), anti-CD19 microbeads (Miltenyi, #130-050-301), anti-CD1c biotin (Miltenyi, #130-113-300) for CD14^+^ and CD19^+^ depletion, as well as with FcR blocking reagent (#130-059-901) in washing buffer (2mM EDTA and 0.1% bovine serum albumin (BSA) in phosphate-buffered saline (PBS)) for 15 minutes at 4 °C. Selected cells were passed through a LD magnetic column (Miltenyi, #130-042-901), followed by a positive selection of CD1c^+^, CD304^+^ and CD141^+^ cells with and anti-biotin microbeads (Miltenyi, #130-090-485), anti-CD304 microbeads (Miltenyi, #130-090-532) and anti-CD141 microbeads (Miltenyi, #130-090-512) that were incubated in washing buffer for 15 minutes at 4 °C. All incubations with microbeads were performed using a ratio of 1 μl/million PBMCs. Cells were then washed and sequentially passed through a LS (Miltenyi, #130-042-401) and MS magnetic column (Miltenyi, #130-042-201). Purity checks were performed (using flow cytometry to ensure the absence of monocyte contamination. In addition, contamination was excluded by gating on CD88^-^ cells (**Supplementary Figure 1A-B**). Experiments were conducted with samples containing >90% purity.

Human pan naïve T cells were isolated using the Pan T cell isolation kit (Miltenyi, #130-097-095535) according to manufacturer’s instructions. Isolated cells were resuspended at 1 × 10⁶ cells/mL in X-VIVO-15 medium (Lonza, #02-053Q) supplemented with 2% human serum (HS, Sigma-Aldrich).

Monocyte-derived dendritic cells (moDCs) were generated from peripheral blood monocytes and differentiated into immature moDCs using IL-4 (500 U/ml, Miltenyi), and GM-CSF (800 U/ml, Miltenyi, #130-093-868). Immature moDCs were then transfected with 3 *LGALS9*-targeting siRNAs (LGALS9HSS142807, LGALS9HSS142808, and LGALS9HSS142809) (Invitrogen)) (or a non-targeting control), and 3 days later matured with IL-6 (15 ng/ml, Miltenyi, #130-093-933), TNF-α (10 ng/mg, Miltenyi, #130-094-014), IL-1β (5 ng/ml, Miltenyi, #130-093-898) and PGE2 (10 µg/ml, Pfizer). Where applicable, *LGALS9* knockdown and DC maturation were monitored by assessing galectin-9 levels and canonical maturation markers as previously reported^26, 27, 30^.

### Dendritic cell and T cell co-culture

DCs were cultured at 37 °C, 5% CO₂ for 48 hours with 50% melanoma CM and 50% X-vivo 2% HS or with X-vivo 2% human serum only at a concentration of 0.5x10^6^ cells/ml at 37 °C, 5% CO₂ for 48 hours. In recovery experiments, DCs were washed after 48 hours of incubation with CM and subsequently incubated for 18 hours in HS-supplemented X-VIVO-15 medium or incubated for 66 hours in CM. When necessary, recombinant human Gal-9/3 protein (1 μg/mL for r-gal-9 and 0,5 μg/mL for r-gal-3, R&D Systems, Minnesota, USA, #2045-GA or #1154-GA-050) was added to CM-treated DCs and incubated for 18 hours in X-VIVO-15 medium supplemented with 2% human serum. r-gal-9 endotoxin content was assessed using Limulus Amebocyte Lysate (LAL) kinetic chromogenic assay performed on a Charles River Endosafe nexgen PTS/MCS system to exclude contamination from the bacterial expression system, and confirmed to be endotoxin-free.

Primary DCs and allogeneic naïve T cells were co-cultured at a ratio of 1:5 (DC:T cell) in a 96 U bottom culture plate. After day 6 (and periodically every two days until day 13-14 of culture), 20 U/ml of recombinant IL-2 (Miltenyi Biotec, #130-097-744) was added to the co-cultures. Expanded T cells were counted and plated for phenotypical analysis.

### Flow cytometry

DCs were detached by adding cold PBS and incubating at 4 °C for 20 minutes to 1 hour. DCs were then incubated with 5% human serum at 4 °C for 10 minutes to block non-specific interactions. and subsequently stained with different surface antibody mixtures (**Table 1**). in cold PBA (PBS containing 0.1% BSA and 0.01% NaN₃) at 4 °C for 30 minutes. Cells were then incubated with an Alexa Fluor 647-conjugated donkey anti-goat secondary antibody in PBA for 20 minutes. For intracellular detection of gal-9 or gal-3, cells were fixed and permeabilized using a FOXP3 transcription factor staining kit (#kit / manufacturer) according to the manufacturer’s instructions prior to being incubated with the corresponding antibody mixture.

**Table 1.** List of antibodies used in this work.

| Application | Antibody Target | Company | Catalog Number | Clone | Color | Dilution |
| --- | --- | --- | --- | --- | --- | --- |
| DC surface gal-9 staining (primary) | Gal-9 | R&D Systems | AF2045 | — | Unconjugated | 5 μg/mL |
| DC surface gal-9 staining (secondary) | Goat IgG (anti-goat) | Thermo Fisher Scientific | A21447 | — | AF 647 | 1:400 (v/v) |
| Human cDC1 | CLEC9A (CD370) | Miltenyi Biotec | 130-106-095 | 8F9 | FITC | 1:40 |
|  | CCR7 (CD197) | BD Biosciences | 560765 | 150503 | PE | 1:25 |
|  | HLA-DR | BioLegend | 307628 | L243 | PerCP | 1:50 |
|  | CD141 (BDCA3) | BioLegend | 344110 | M80 | PE-Cy7 | 1:25 |
|  | CD80 | BD Biosciences | 561133 | L307.4 | AF 700 | 1:25 |
|  | CD86 | BD Biosciences | 561124 | 2331 (FUN-1) | AF 700 | 1:25 |
|  | PD-L1 (CD274) | BD Biosciences | 563738 | MIH1 | BV421 | 1:25 |
|  | CD83 | BioLegend | 305338 | HB15e | BV785 / BV786 | 1:25 |
| Human cDC2 and CD14<sup>+</sup> acquisition | CD88 (C5aR) | BioLegend | 344304 | S5/1 | PE | 1:25 |
|  | HLA-DR | BioLegend | 307628 | L243 | PerCP | 1:50 |
|  | PD-L1 (CD274) | BD Biosciences | 558017 | MIH1 | PE-Cy7 | 1:25 |
|  | CCR7 (CD197) | Miltenyi Biotec | 130-120-466 | REA546 | APC | 1:25 |
|  | CD80 | BD Biosciences | 561133 | L307.4 | AF 700 | 1:25 |
|  | CD1c (BDCA1) | BioLegend | 331526 | L161 | BV421 | 1:25 |
|  | CD5 | BioLegend | 300644 | UCHT2 | BV711 | 1:25 |
|  | CD14 | BD Biosciences | 563698 | M5E2 | BV786 | 1:25 |
| Human pDCs | CD303 (BDCA2) | Miltenyi Biotec | 130-113-193 | AC144 | PE | 1:25 |
|  | CD40 | BD Biosciences | 561215 | 5C3 | PE-Cy7 (PE-Vio770) | 1:25 |
|  | CD123 (IL-3Rα) | BD Biosciences | 663994 | 9F5 | APC-R700 | 1:25 |
|  | CD123 (IL-3Rα) | BioLegend | 306039* | 6H6 | AF 700 | 1:25 |
|  | OX40L (CD252) | BD Biosciences | 563766 | ik-1 | BV421 | 1:25 |
|  | HLA-DR | BioLegend | 307646 | L243 | BV510 | 1:25 |
|  | CD83 | BioLegend | 305338 | HB15e | BV785 / BV786 | 1:25 |
| Human T cell exhaustion | CD4 | BD Biosciences | 564975 | RPA-T4 | APC-R700 | 1:50 |
|  | CD8 | BD Biosciences | 555366 | RPA-T8 | FITC | 1:100 |
|  | TIGIT | BD Biosciences | 747846 | 741182 | BV786 | 1:20 |
|  | Tim-3 | BioLegend | 119703 | RMT3-23 | PE | 1:100 |
|  | LAG-3 | Thermo Fisher Scientific | 63-2239-42 | 3DS223H | Super Bright 600 | 1:10 |
|  | OX40 | BioLegend | 119413 | OX-86 | APC | 1:10 |
|  | CTLA-4 | BD Biosciences | 555853 | BNI3 | PE | 1:20 |
|  | ICOS | BD Biosciences | 562833 | DX29 | PerCP-Cy5.5 | 1:20 |
|  | PD-1 | BioLegend | 329919 | EH12.1 | BV421 | 1:15 |
| Human CD4+-cytokine production | IL-10 | BioLegend | # 501420 | JES3-9D7 | PE-Cy7 | 1:20 |
|  | IL-4 | Miltenyi Biotec | # 130-123-698 | 7A3-3 | PE | 1:25 |
|  | IFN-γ | Miltenyi Biotec | # 130-113-497 | REA600 | FITC | 1:50 |
|  | IL-17A | BioLegend | # 512310 | BL168 | AF647 | 1:25 |
|  | CD4 | BioLegend | # 300528 | RPA-T4 | PerCP | 1:25 |
| Human CD8+-granular production | Granzyme-B | BioLegend | # 372208 | QA16A02 | PE | 1:30 |
|  | Perforin | BioLegend | # 308110 | dG9 | af647 | 1:30 |
|  | CD8 | BD Biosciences | # 555366 | RPA-T8 | FITC | 1:100 |
| Human T regulatory phenotype | CD4 | eBioscience | # 25-0047-42 | SK3 | PE-Cy7 | 1:50 |
|  | CD25 | BD Biosciences | # 555434 | M-A251 | APC | 1:100 |
|  | Foxp3 | eBioscience | # 53-4776-42 | PCH101 | af488 | 1:50 |
|  | CD127 | eBioscience | # 12-1278-42 | eBioRDR5 | PE | 1:100 |
| Human CD4+-T helper subsets | CD4 | eBioscience | # 12-1278-42 | eBioRDR5 | PE | 1:50 |
|  | GATA3 | BD Biosciences | # 560078 | L50-823 (RUO) | af647 | 1:100 |
|  | T-bet | Invitrogen | # 45-5825-82 | 4B10 | PerCP-Cy5.5 | 1:100 |
|  | RORγt | Invitrogen | # 12-6988-82 | AFKJS-9 | PE | 1:200 |
| DC glycome (lectin binding) | LEL (biotinylated) | Vector Labs | B-1175-1 | — | Biotinylated lectin | 10 µg/mL |
|  | SNA-I (biotinylated) | Vector Labs | B-1305-2 | — | Biotinylated lectin | 10 µg/mL |
|  | UEA-I (biotinylated) | Vector Labs | B-1065-2 | — | Biotinylated lectin | 10 µg/mL |
|  | LCA (biotinylated) | Vector Labs | B-1045-5 | — | Biotinylated lectin | 10 µg/mL |
|  | DBA (biotinylated) | Vector Labs | B-1035-5 | — | Biotinylated lectin | 10 µg/mL |
|  | PHA-L (biotinylated) | Vector Labs | B-1115-2 | — | Biotinylated lectin | 10 µg/mL |
| Lectin detection | Streptavidin | BioLegend | 405213 | — | PerCP-conjugated | 1:1000 (v/v) |
|  | Streptavidin | BioLegend | 405201 | — | FITC-conjugated | 1:1000 (v/v) |

To determine glycan changes cells were incubated with the corresponding biotinylated lectin (LEL, SNA-I, UEA-I, LCA, DBA and PHA-L) at 4 °C for 1 hour in lectin buffer (Carbo-Free Blocking Solution (VectorLabs, #SP-5040-125) supplemented with 1 mM NaCl and 1 mM MgCl) followed by incubation with PerCP- or FITC-conjugated streptavidin in lectin buffer for 30 minutes at 4 °C.

To determine T cell immunophenotypes, harvested T cells were blocked in PBA supplemented with 2% human serum for 10 min and then divided into different flow-cytometry panels. Exhausted T cells were identified by surface staining for CD4, CD8, TIGIT, Tim-3, LAG-3, OX40, CTLA-4, ICOS and PD-1 (**see Table 1** for antibody details). To determine regulatory T cell subsets, T cells were fixed and permeabilized with the FOXP3 transcription factor staining kit and blocking as described above. Regulatory T cell subsets were identified by staining for CD4, CD25, FOXP3 and CD127, whereas T-helper subsets were defined using antibodies against CD4, GATA3, T-bet and RORγt (**Table 1**).

To determine intracellular cytokine content, CD4⁺ and CD8⁺ cells were stimulated with PMA (25 ng/mL), ionomycin (0.5 μg/ml) and brefeldin A (10 ng/ml) for 4 hours at 37 °C. Cells were washed and live/dead staining was performed with eFluor780 or zombie violet viability dyes (1:2,000, ThermoFisher) in PBS for 20 min at 4 °C, then fixed and permeabilized using the BD Cytofix/Cytoperm™ Fixation/Permeabilization Kit according to the manufacturer’s instructions. After blocking, cells were stained with antibodies against IL-10, IL-4, IFN-γ, IL-17A and CD4 for analysis of CD4⁺ T cells, and with antibodies against Granzyme B, Perforin and CD8 for analysis of CD8⁺ T cell granule content (**Table 1**).

Live/dead stainings were performed by incubating cells with eFluor780 or zombie violet viability dyes (1:2000, ThermoFisher) in PBS for 20 min at 4 °C prior to being fixed with 2% paraformaldehyde in PBA at room temperature for 5 minutes. Samples were stored at 4 °C until acquisition. Samples were acquired by flow cytometry using a FACSVerse or a FACSLyric instrument (BD) and later analyzed using the Cytobank platform (Beckman Coulter Life Sciences)^31^.

### RNA isolation and sequencing

Human primary DCs pellets (treated with or without CM) were processed immediately for RNA extraction. A minimum amount of 0.25x10^6^ cells per condition was used. Total RNA was isolated using Quick-RNA™ Miniprep Kit (Zymo Research) according to the manufacturer’s instructions. Eluted RNA was quantified and stored at -80°C until analysis. RNA purity was assessed by A260/A280 and A260/A230 ratios.

RNA-seq libraries were prepared using 50ng input RNA per sample, bulk RNA barcoding and sequencing (BRB-seq^32^) (Alithea Genomics) based on the manufacturer’s instructions. The pooled library was sequenced on a Nextseq 2000 (Illumina). Reads were aligned to the hg38 reference human genome and demultiplexed to individual samples based on the barcode sequences using STARsolo^33^. Differential gene expression was calculated using DEseq2^34^. Gene ontology enrichment was performed using EnrichR^35^. All downstream analyses were performed using R.

### Single-cell RNA sequencing analysis

Single cell RNA-seq data for tumor-associated myeloid-derived cells and healthy PBMCs were generated previously^36^, and downloaded from the CELLxGENE platform^37, 38^. The original integration and embedding were used. DCs from tumor sites and healthy PBMCs were subsetted for visualization. Myeloid-derived cells were analyzed by integrating multi-cancer single-cell RNA sequencing datasets through a standardized pipeline involving Harmony-based batch correction, Seurat-mediated clustering for subpopulation identification, and prognostic data extraction via deconvolution of bulk transcriptomic cohorts.

### Statistical analysis

All data was processed using Excel (Microsoft), analyzed and plotted using GraphPad Prism (Version 8.0.2. or 10.0.0 GraphPad Software). All data is expressed as mean +/-SEM. The statistical test used to analyze each data set is described in the corresponding figure legend. Data distribution was assessed using the Shapiro-Wilk’s test. Statistical significance threshold was defined as ∗p < 0.05; ∗∗p < 0.05; ∗∗∗p < 0.005.

## Results

### Tumor exposure reduces gal-9 surface levels in immunosuppressed DCs

Contrary to other tumors, in melanoma gal-9 is predominantly anti-tumorigenic, as higher gal-9 expression has been associated with increased melanoma cell apoptosis, reduced metastasis, and improved patient survival^39, 40, 41, 42^. Studying gal-9 in dendritic cells is particularly relevant in melanoma, where DC-expressed gal-9 has been associated with improved clinical outcomes and may contribute to the regulation of anti-tumor immunity^43^. To examine whether gal-9 expression in DCs changed upon exposure to tumors, we isolated naïve cDC2 DCs from healthy PBMCs, incubated them with a panel of melanoma-derived conditioned media (CM) for 48 or 72 hours after which gal-9 expression was determined by flow cytometry. Exposing DCs to melanoma CM resulted in a loss of surface gal-9 for all cell lines used, being the A375 cell line the one provoking a greater decrease (**Supplementary Figure 1B-C**).

Next, we determined whether changes in surface gal-9 expression were specific to DC2s or also occurred in other DC subsets. To meet this end, we isolated primary human cDC1, cDC2, CD14^+^cDC2, and pDCs from healthy PBMCs and cultured them for 48 hours with A375-derived-CM (**Figure 1** and **Supplementary Figure 2**). Following exposure to tumor-associated factors, cDC1s displayed reduced surface HLA-DR, CD80, PDL1 and CCR7 expression, alongside a slight increase in CD83 levels, consistent with the acquisition of an immunosuppressed phenotype (**Supplementary Figure 2B**). cDC1 were found to express very little gal-9, which decreased to undetectable levels upon culturing with melanoma CM (**Supplementary Figure 2B**). pDCs showed no significant changes in HLA-DR and CD83 surface expression but displayed elevated CD40, suggesting a mild activation driven by the inflammatory components of the CM (**Supplementary Figure 2D**). As observed for cDC1, pDCs displayed undetectable levels of gal-9 under naïve conditions as well as upon exposure to CM (**Supplementary Figure 2D**). Upon melanoma CM-treatment, cDC2s acquired CD14 expression (>50% of cells), generating a heterogeneous population of CD1c^+^CD14^-^ and CD1c^+^CD14^+^ (hereby referred as cDC2 and CD14^+^cDC2) cells, as previously reported^6, 7^ (**Supplementary Figure 1D**). cDC2s showed no significant changes in CD80, gal-9 or PDL1 but displayed enhanced HLA-DR and CCR7 surface expression, suggesting the acquisition of an activated phenotype driven by the CM inflammatory properties (**Figure 1B**). In contrast, CD14^+^cDC2 cells exhibited immunosuppressive features, evidenced by the downregulation of HLA-DR and upregulation of PDL1, whilst CCR7 and CD80 remained unaltered (**Figure 1C**). Notably, and contrary to all other DC subsets, CD14^+^cDC2 cells showed a significant reduction in gal-9 levels after melanoma-CM exposure, indicating that gal-9 loss in DCs within the TME is associated to specific DC subsets.

**Figure 1.**
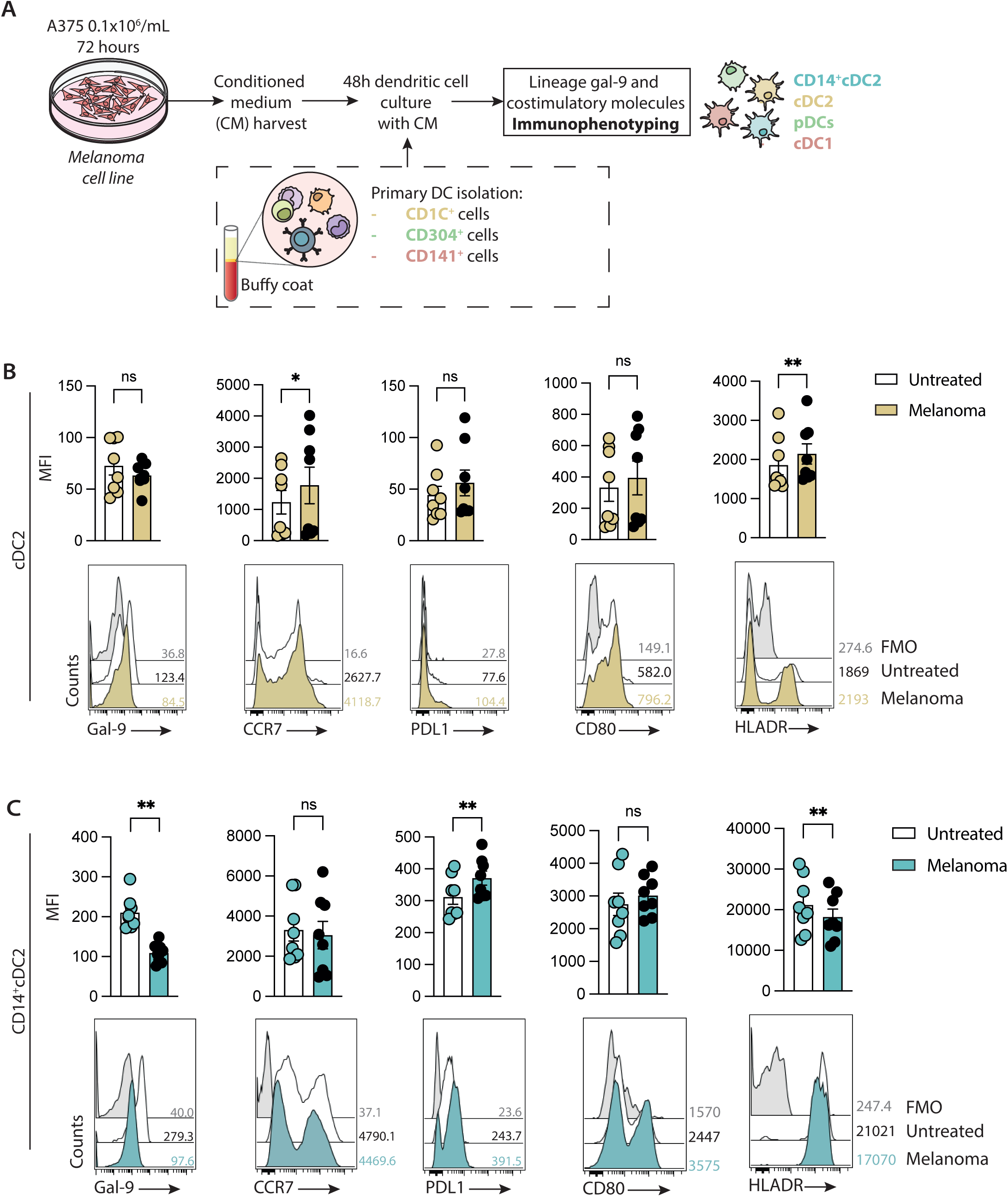
Melanoma-conditioned media (CM)-derived CD14^+^cDC2s exhibit reduced gal-9 levels. **A.** Schematic representation of experimental layout: A375 melanoma cell line was cultured at 0.1x10^6^ cells/mL during 72 hours and the media was harvested and used to culture human primary DCs for 48 hours. Lineage and activating or inhibitory receptors for each cell type were immunophenotyped using flow cytometry. **B and C.** Representative flow cytometry histogram (down) and mean fluorescent intensity (MFI) quantification (up) (n=8) for Gal-9, CCR7, PDL1, CD80 and HLA-DR markers. Unstained control or fluorescence minus one (FMO) samples are shown in gray, untreated samples are displayed in white and melanoma-CM treated samples in yellow (for cDC2) or blue (for CD14^+^cDC2s). Numbers in histograms indicate the MFI value of each histogram. Data shown as mean ± SEM. Each dot represents an independent donor. Statistical significance assessed by paired student T test. ns p > 0.05, *p < 0.05 and **p < 0.01.

### Tumor-derived high-molecular-weight factors induce surface proteome loss of gal-9 in CD14^+^cDC2s

To gain deeper insights into the molecular mechanisms underlying the decreased levels of surface gal-9 in immunosuppressed cDC2s cells, we first sought to investigate whether this reduction also occurred intracellularly. Analysis of gal-9 levels in permeabilized CD14^+^cDC2s revealed that intracellular abundance of this lectin remained high in DC2s exposed to CM (**Figure 2A** and **2B**). Next, we assessed whether the decrease in surface gal-9 in cDC2s was reversible upon removal of tumor pressure. To this end, cDC2s cultured with melanoma CM were washed, incubated in normal media for 18 hours (hereby referred to as recovery), and their gal-9 levels were compared to cells continuously exposed to CM (hereby referred to as non-recovery) (**Figure 2C**). Cells under constant CM exposure maintained reduced surface gal-9 levels, whereas those relieved from tumor factors restored gal-9 to baseline levels, indicating that this regulation is transient and likely not transcriptionally driven (**Figure 2D** and **2E**). To confirm this, we performed RNA sequencing from DCs treated or not with CM. Validating our approach, genes previously described to increase in DCs after tumor encounter were also enhanced in our system (**Figure 2F**)^44^. Importantly, genes involved in antigen presentation, immunosuppression programs, CD14^+^cDC2 identity, metabolism and membrane organization were upregulated upon melanoma treatment compared to untreated samples (**Supplementary Figure 3A-B**). As shown, gal-9 transcript numbers (*LGALS9*) did not show any significant differences before and after melanoma treatment (**Figure 2F**). To confirm this result in intra-tumoral DCs, we analyzed publicly available single-cell RNA sequencing (sc-RNA-seq) datasets PBMCs exposed or not (healthy) to melanoma (**Figure 2G**)^36^. From an unbiased clustering analysis, we obtained *LGALS9* expression in cDC2s and in CD14^+^cDC2 (hereby annotated as cDC2_FCER1A and cDC3_CD14, respectively) in DCs resident in melanoma compared to healthy PBMCs. We observed no significant differences in *LGALS9* expression between these subsets, supporting that gal-9 differential expression upon CM exposure occurs at the protein level rather than transcriptomically (**Figure 2H** and **Supplementary Figure 3C**). These findings were validated in a murine system using sc-RNA-seq data from B16 melanoma tumors at days 5, 8, and 11 after subcutaneous injection (**Supplementary Figure 3D-F**). *Lgals9* was not altered transcriptionally over time in cDC1, cDC2/3 or pDCs (**Supplementary Figure 3G**).

**Figure 2.**
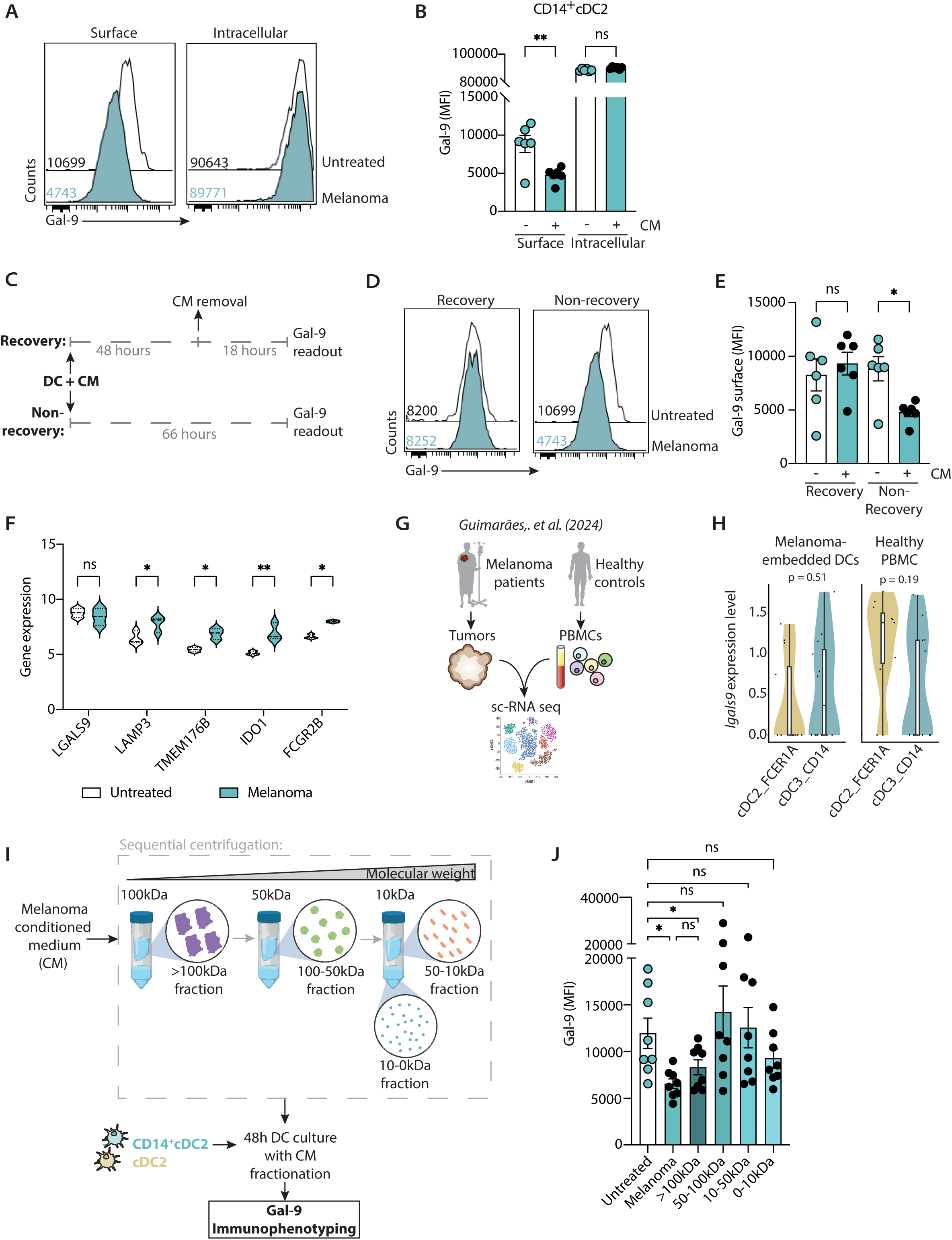
Melanoma >100kDa components induce a reduction of surface gal-9 in CD14^+^cDC2s. **A.** Representative flow cytometry histogram of the surface (left) or intracellular (right) expression of gal-9 in untreated (white) or melanoma-exposed CD14^+^cDC2s (blue). **B.** Quantification of gal-9 MFI (n=6) of A. **C.** Schematic diagram of the experimental timeline; CD1c^+^ cells were incubated for 48 hours with melanoma CM, washed and left with normal media for 18 remaining hours (recovery) or exposed with melanoma CM for 66 hours (non-recovery). In both cases, expression of gal-9 was measured by flow cytometry. **D.** Representative flow cytometry histogram of gal-9 expression after the recovery (left) or non-recovery (right) conditions in untreated (white) or melanoma-exposed CD14^+^cDC2s (blue). **E.** Quantification of gal-9 MFI (n=6) of D. **F.** Violin plots showing RNA-seq data obtained from melanoma-CM exposed vs untreated DCs isolated RNA. Gene (*IDO1, LAMP3, TMEM176B, LGALS9,* and *FCGR2B*) expression levels is shown in white for untreated and blue for melanoma-exposed DCs. **G.** Schematic overview of the experimental design adapted from Guimarães et al. (2024). sc-RNA seq was performed in human melanomas and healthy PBMC controls to evaluate *lgals9* expression in specific DC subsets. **H.** Violin plots showing *lgals9* expression levels in cDC2 (CDC2_FCER1A, in yellow) and CD14^+^cDC2 (CDC3_CD14 in blue) in samples obtained from melanoma patients (left) or healthy controls (right). Statistical comparisons between signatures were performed using a Wilcoxon rank-sum test. **I.** Sequential fractionation of melanoma-CM and DC culture workflow. Melanoma-CM was separated by molecular weight using sequential centrifugation into four fractions: >100 kDa, 100-50 kDa, 50-10 kDa, and 10-0 kDa. CD14⁺ cDC2s and cDC2 subsets were cultured for 48 hours with CM fractions, followed by gal-9 immunophenotyping. **J.** Quantification of gal-9 MFI (n=8) of untreated (white) or fractionated CM-exposed CD14^+^cDC2s (gradients of blue). Numbers in histograms indicate the MFI value of each histogram. Data shown as mean ± SEM. Each dot represents an independent donor. Statistical significance assessed by two-way ANOVA with Šídák’s multiple comparisons or a Wilcoxon rank-sum test (in H). ns p > 0.05, *p < 0.05 and **p < 0.01.

Next and to gain further insights into the tumor-derived components responsible for the reduction in gal-9, we fractionated CM into distinct molecular weight fractions (>100 kDa, 100-50 kDa, 50-10 kDa, and <10 kDa) (**Figure 2I**). Treating CD14^+^cDC2 cells with the >100kDa fraction recapitulated the loss of surface gal-9 observed upon treatment with full melanoma-CM, whereas gal-9 levels remained unaltered upon incubation with all other fractions (**Figure 2J**). Gal-9 did not decrease in cDC2s upon incubation with any CM molecular fraction, substantiating the specificity of gal-9 loss in immunosuppressed CD14^+^cDC2 cells, and implicating high-molecular-weight compounds as drivers of this effect (**Supplementary Figure 4A**).

Finally, we assessed the specificity of the phenotype within the galectin family by examining galectin-3 (gal-3) expression in CD14⁺cDC2s upon melanoma-CM exposure. Gal-3 intracellular and surface levels were not affected in response to melanoma-CM pressure (**Supplementary Figure 4B).** Moreover, no changes in gal-3 expression were observed upon removal of melanoma-CM and the >100kDa CM fraction did not elicit any changes in gal-3 compared to untreated conditions; interestingly, gal-3 levels were shown to decrease upon incubation with 0-10 kDa fraction (**Supplementary Figure 4C-D**).

Altogether, our data confirm that CD14⁺cDC2s specifically decrease cell surface gal-9 and suggest the existence of distinct regulatory mechanisms underlying expression of different members of this lectin family.

### Gal-9 loss is accompanied by changes in the DC glycome

Given that gal-9 loss upon exposure to CM was not associated with transcriptional changes, we next investigated whether it might depend on the repertoire and structure of cell surface glycans and the availability of gal-9 ligands. To that end, we examined the DC glycome before and after tumor exposure using fluorescent streptavidin-conjugated plant lectins (**Figure 3A**)^23^. In addition, we also compared naïve CD14⁺ and CD14⁻ cDC2 subsets to determine whether the CD14⁺ population displays a distinct glycan profile from its CD14⁻ counterpart and whether melanoma-CM induces unique glycan signatures in each population. Both CD14⁻ cDC2 and CD14⁺ cDC2s displayed high abundance of β(1,6)-branched N-glycans (PHA-L) and mannose residues (LCA), which remained unchanged after melanoma CM treatment (**Figure 3B-E**). Analysis of the relative composition of each glycoepitope in relation to the others (in %) across 5 independent donors revealed that CD14⁻ cDC2s and CD14⁺ cDC2s display similar glycophenotypes under naïve conditions (untreated) (**Figure 3C**). Upon exposure to melanoma CM, both subsets showed significantly reduced DBA binding, which can be associated to lower α-linked N-acetylgalactosamine (GalNAc) and particularly blood group A structures. In addition, α(1,2) fucosylation (shown by UEA-I labeling) was reduced only in CD14⁻ cDC2s whereas a2,6-sialylation (evidenced by SNA-I binding) was lower only in CD14⁺cDC2s (**Figure 3D-E**).

**Figure 3.**
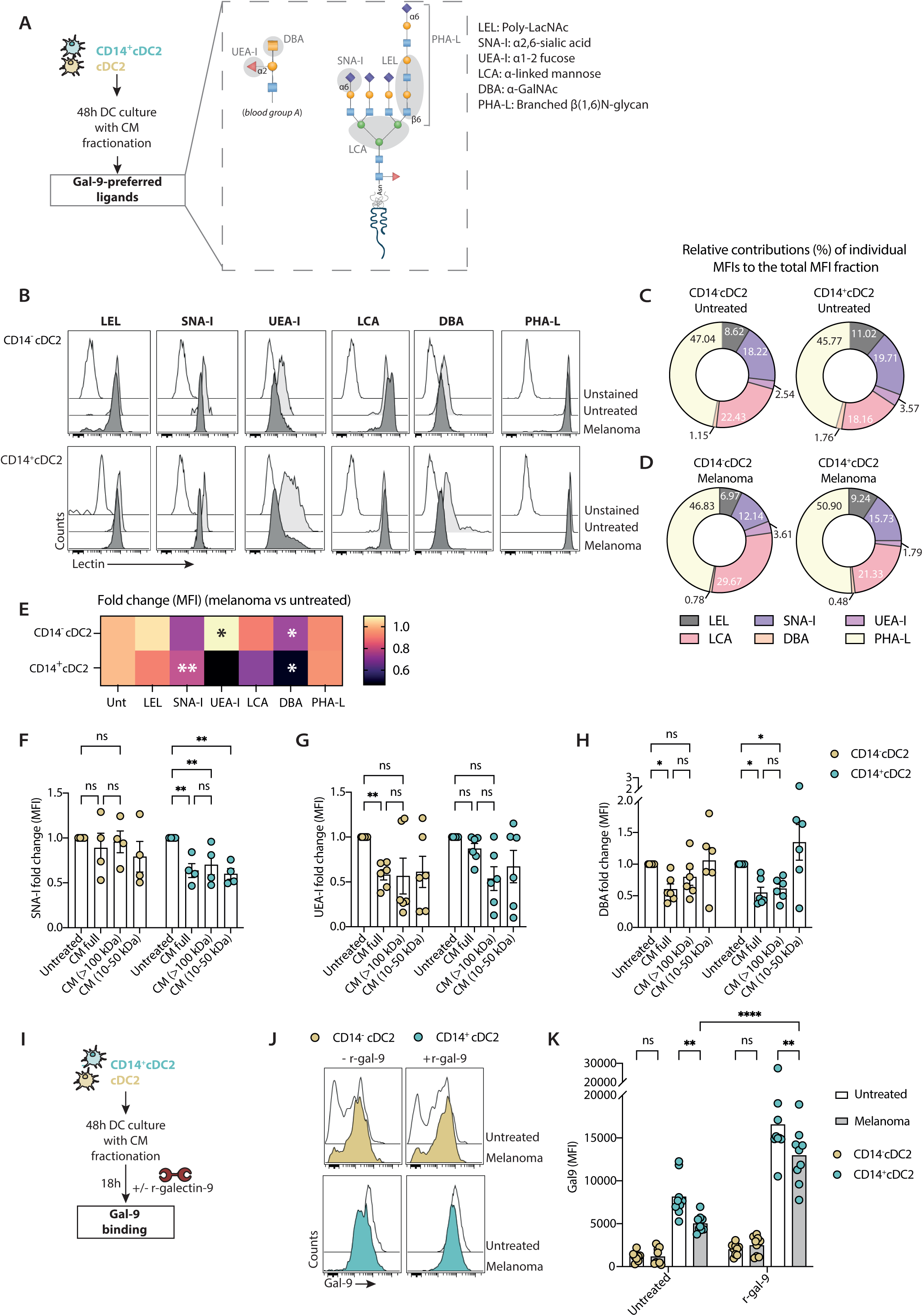
Glycan profiling of cDC2 and CD14⁺cDC2 subsets under untreated and melanoma-conditioned conditions. **A.** CD14⁺ cDC2 and cDC2 subsets were cultured for 48 hours with melanoma-CM fractions, and glyco-binding residues susceptible to gal-9 interaction were analyzed by flow cytometry. The schematic shows representative glycan motifs, including blood group antigen A and branched N-glycans, with lectin-binding sites indicated in gray (LEL: poly-lacNAc; SNA-I: α2,6-sialic acid; UEA-I: α1,2 fucose; DBA: GalNAc; LCA: α-linked mannose; PHA-L: branched N-glycans). **B.** Representative flow cytometry histograms showing lectin binding to cDC2 (top row) and CD14⁺cDC2 (bottom row) subsets. White histograms represent unstained controls, gray histograms represent untreated cells, and dark-gray histograms represent melanoma-CM-treated cells. **C-D.** Donut charts summarizing the proportion of each MFI (in %) within the total MFI fraction on CD14^-^cDC2 and CD14⁺cDC2 subsets in untreated (**C**) and melanoma-conditioned (**D**) conditions. **E.** Heat map showing normalized melanoma-treated cells lectin binding intensities to untreated, statistical significance is represented with a white * (n=5). **F-H.** Fold change relative to untreated fom the quantification of lectin binding (MFI) for DBA (F), UEA-I (G) and SNA-I (H) across conditions: untreated, melanoma-conditioned, and fractions from melanoma-CM. **I.** Experimental layout depicting the r-gal-9 addition to CM-treated DCs timeline. **J.** Representative flow cytometry histograms showing gal-9 expression (MFI) in cDC2s and CD14⁺ cDC2s. Cells were analyzed under untreated conditions or following exposure to melanoma-CM, in the absence (- r-gal-9) or presence (+ r-gal-9) of recombinant gal-9. **K.** Quantification of I (n=9). Yellow dots represent cDC2s and blue dots CD14⁺ cDC2s. Data shown as mean ± SEM. Each dot represents an independent donor. Statistical significance assessed by two-way ANOVA with Šídák’s multiple comparisons. ns p > 0.05, *p < 0.05, **p < 0.01 and ***p < 0.001.

To further explore the correlation between gal-9 loss and the altered DC glycophenotype upon CM treatment, we further analyzed SNA-I, DBA and UEA-I lectin binding at the DC surface after exposing CD14⁻ and CD14^+^ DC2s to differential molecular weights CM fractions. All CM fractions reduced SNA-I binding compared to untreated cells in CD14⁺ cDC2s and the same pattern was observed for UEA-I in CD14⁻ cDC2s (**Figure 3F-G**). Interestingly, α-GalNAc levels (DBA binding) were significantly decreased in CD14^+^ cells upon treatment with either full CM or the >100 kDa CM fraction, mirroring the pattern of gal-9 reduction previously observed (**Figure 3H**). Given that gal-9 displays preferential binding to blood group like antigens^21^, these findings support a model in which high-molecular weight melanoma-derived factors might induce a specific glycan remodeling in DCs, thereby resulting in the loss of gal-9 ligands, particularly GalNAc-contaning blood group structures at the DC surface.

To further confirm that melanoma-CM-induced remodeling of the DC glycome modulates gal-9 binding, we treated DCs with recombinant gal-9 (r-gal-9) in the presence or absence of CM (**Figure 3I-J**). R-gal-9 exhibited a lower binding capacity to CD14^-^ cDC2s cells compared with their immunosuppressed CD14⁺ counterparts already in a naïve state, indicating differential gal-9 binding or glycan expression between the two subsets. In addition, a reduction in r-gal-9 binding was observed in DC2s exposed to melanoma CM, most notably in CD14^+^ cells (**Figure 3K**), consistent with melanoma-induced glycan remodeling on the DC surface, and subsequently hindering gal-9 recognition and binding.

### Gal-9 rescue impedes immunosuppressed DC-mediated Treg expansion

Tumor-conditioned DCs can impose T cell-dependent tolerogenic circuits, most prominently by expanding Tregs, promoting dysfunctional/exhausted states, and weakening effector differentiation, thereby supporting tumor progression^45^. Thus, we next asked whether loss of gal-9 on CD14⁺ cDC2s contributes to altered T cell polarization. To address this, we co-cultured cDC2s pre-treated with CM, and in the presence or absence of r-gal-9, with naïve pan T cells and assessed effector and regulatory populations by flow cytometry (**Figure 4A and Supplementary Figure 5**).

**Figure 4.**
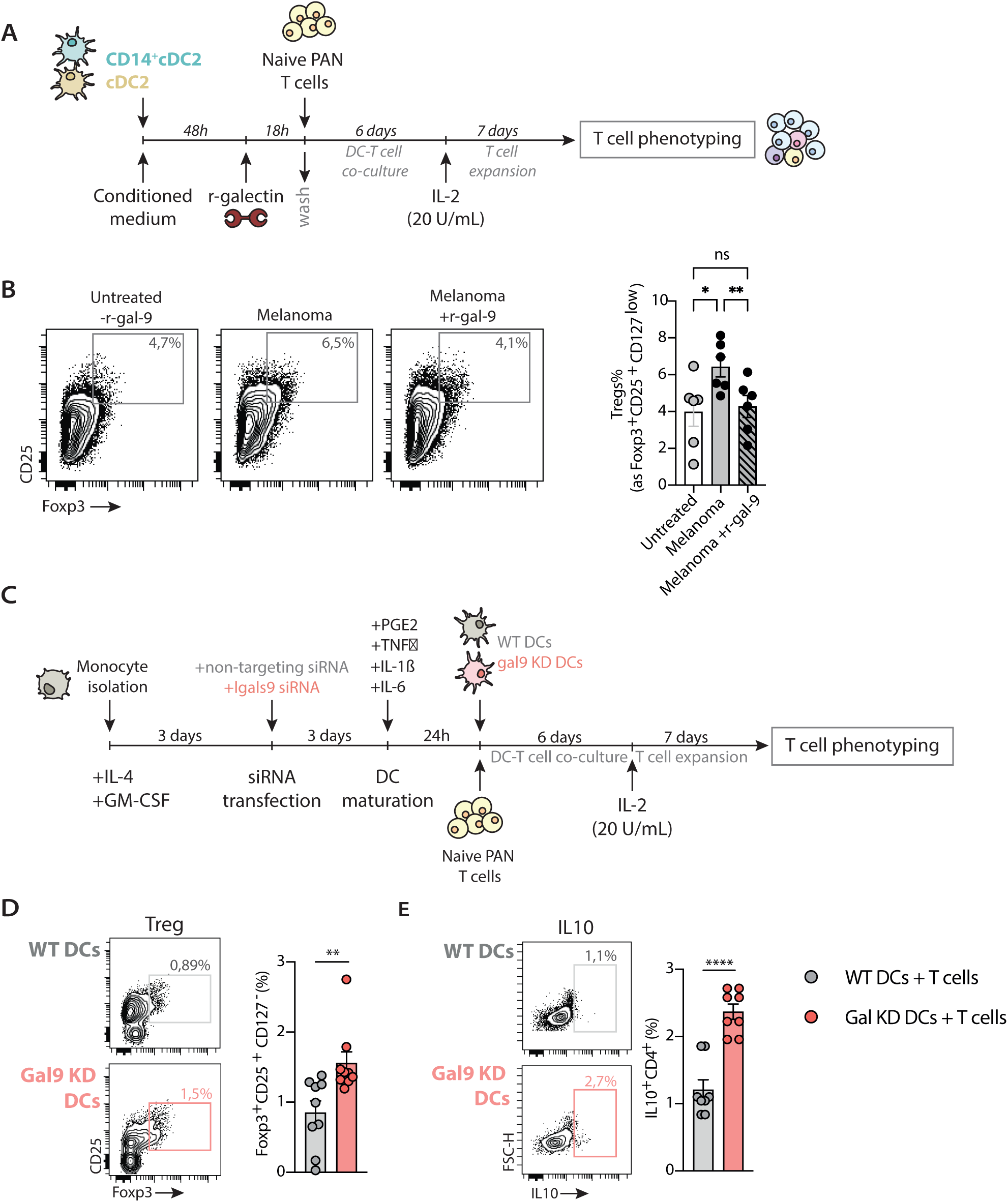
Functional T cell polarization studies show that restoring gal-9 in immunosuppressed DCs restores Treg frequencies and effector functions to untreated levels. **A.** Experimental workflow for DC-T cell co-culture assays. DCs were exposed to melanoma-CM and/or recombinant gal-9, followed by 6-day co-culture with naïve pan T cells and a T cell expansion phase in the presence of IL-2 (20 U/mL). T cell phenotyping was performed after 7 days using flow cytometry. **B.** Representative flow cytometry plots (left) and quantification of Tregs (CD25⁺CD127^low^Foxp3⁺) (right) (n=6) generated in co-cultures with untreated or melanoma-conditioned DCs ± gal-9. **C.** Schematic of siRNA-mediated gal-9 knockdown (KD) in DCs followed by maturation and co-culture with naïve PAN T cells. **D.** Effect of gal-9 KD on moDC-mediated Treg induction. Representative flow contour plots (left) and quantification (right) (n=9) comparing wildtype (WT) DCs (gray) and gal-9 KD DCs (red). **E.** Effect of gal-9 KD on moDC-mediated IL-10-producing cells induction. Representative flow contour plots (left) and quantification (right) (n=8) comparing WT DCs (gray) and gal-9 KD DCs (red). Data shown as mean ± SEM. Each dot represents an independent donor. Statistical significance assessed by two-way ANOVA with Šídák’s multiple comparisons. ns p > 0.05, *p < 0.05, **p < 0.01, ***p < 0.001 and ****p < 0.0001.

Following r-gal-9 treatment, DCs exposed or not to melanoma CM showed no significant differences in inducing Th1, Th17, and Th2 responses, as evidenced by increased IFN-γ (Th1) and IL-17 (Th17) as well as diminished IL-4 (Th2) (**Supplementary Figure 6A**). Melanoma exposed CD14^+^cDC2 induced Treg polarization, a phenotype which could be reverted by treating DCs with r-gal-9 prior to being co-cultured with T cells (**Figure 6B**). Assessment of T cell exhaustion revealed that melanoma-CM increased exhaustion markers, but r-gal-9 treatment did not significantly restore these phenotypes, suggesting that gal-9 in DCs specifically modulates Treg polarization (**Supplementary Figures 5D, 6B**). Interestingly, treating DCs with recombinant gal-3 (r-gal-3) did not alter Treg polarization, suggesting a specific role for gal-9 in T cell differentiation (**Supplementary Figure 6C**).

Overall, these findings suggest that tumor-induced CD14^+^cDC2s preferentially prime Treg and Th2 responses while dampening Th1/Th17 polarization, and r-gal-9 treatment mitigates these effects mostly in the Treg population.

To validate a role for gal-9 in DC-mediated T cell priming, we performed *LGALS9* knockdown in monocyte-derived DCs and co-cultured these with naïve T cells (**Figure 4C**). moDCs devoid of gal-9 did not induce changes in T helper subset transcription factors (T-bet, RORγt, GATA3), cytokine secretion (IFN-γ, IL-17, IL-4), exhaustion or CD8^+^ cytolytic activity (**Supplementary Figures 5A, C-F and 7A-D**). However, gal-9-deficient moDCs induced higher Treg frequencies and increased IL-10 secretion compared to their WT counterparts (**Figures 4D-E**), confirming that gal-9 plays a protective role in limiting Treg induction and maintaining DC immunostimulatory function.

## Discussion

In melanoma, gal-9^high^-expressing DCs were correlated with positive long-term clinical outcomes^43^. DC-mediated anti-melanoma immunity has been mechanistically explored mainly by transcriptomic and proteomic analyses, leaving tumor-induced glycosylation changes in DCs and their functional impact largely underexplored. In this study we demonstrate how melanoma-derived soluble factors selectively and transiently reduce surface gal-9 specifically on immune suppressed DCs (CD14⁺cDC2) via a transcription-independent mechanism. Fractionation of the melanoma secretome revealed that >100 kDa components drive this effect and remodel the CD14⁺ cDC2 glycome towards a non-permissive gal-9-binding profile. In addition, CD14+ DC2 displayed impaired Treg expansion, whereas restoring surface gal-9 rescued Treg induction to levels seen in non-tumor-exposed DCs. Thus, we hypothesize that high-molecular weight melanoma-derived factors reprogram the CD14⁺ cDC2 glycome, in turn diminishing surface gal-9 binding, and enhancing DC-mediated Treg induction (**Figure 5**). To our knowledge, this is the first evidence of tumor-induced glycome alterations in DCs with functional consequences for gal-9 binding and Treg polarization.

**Figure 5.**
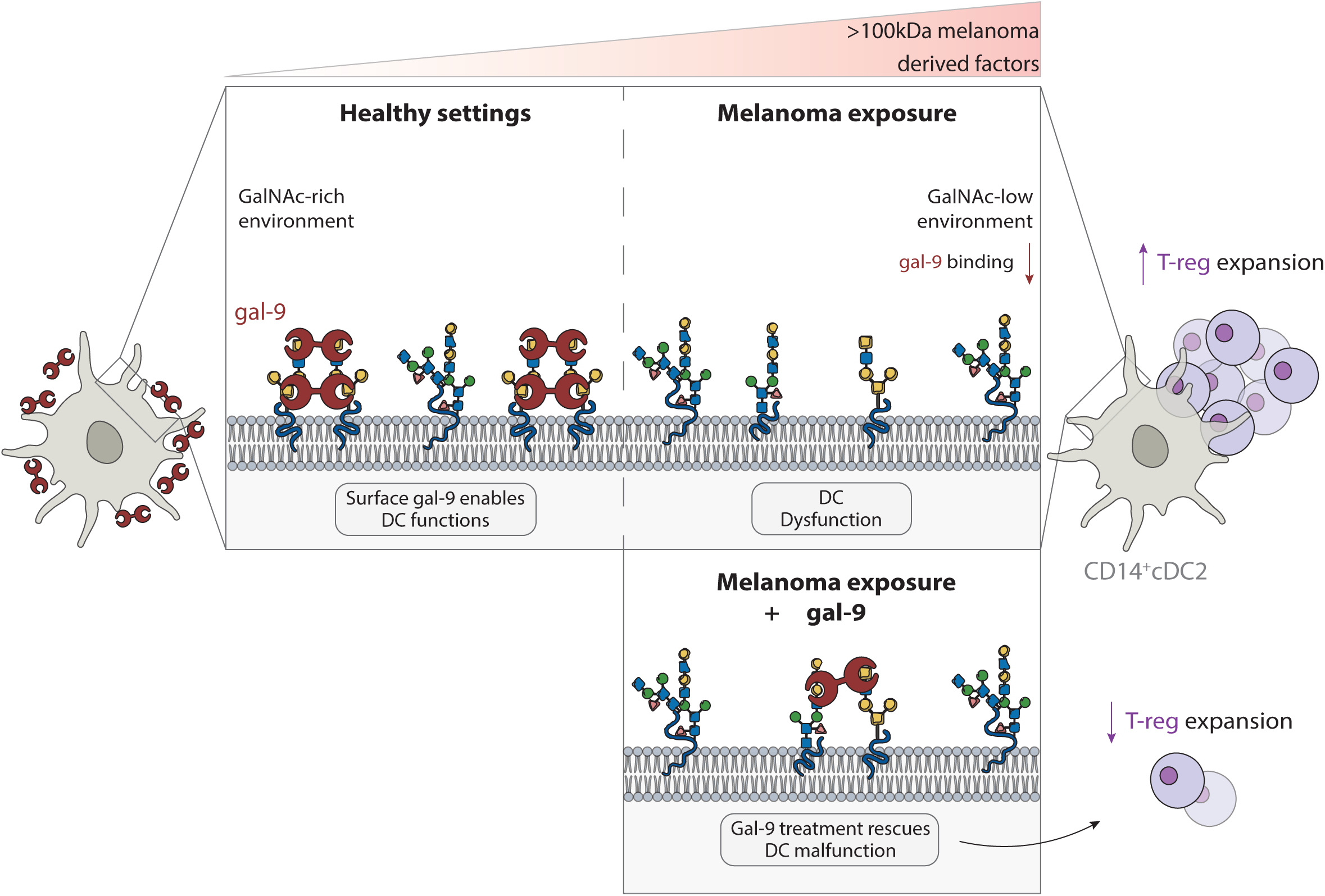
Graphical abstract. Proposed model illustrating how melanoma affects gal-9 expression and binding capacity on the surface of DCs. In healthy settings, CD14^+^cDC2 display high levels of surface gal-9 which correlates with GalNAc residue abundance and supports proper DC function. Upon exposure to >100 kDa melanoma-derived factors, gal-9 binding to DC surface is reduced, correlating with less GalNAc residues and leading to DC dysfunction and functional outcomes such as expansion of T regulatory (Treg) populations. Supplementation with r-gal-9 shows reduced binding capacity but functionally rescues DC activity, thereby limiting Treg expansion.

Our finding that CD14^+^cDC2s display lower gal-9 levels after exposure to melanoma-CM is consistent with a wide range of studies that reported altered galectin expression in human malignancies, often correlating with tumor progression and patient outcome^32^. In particular, gal-9 expression shows tumor-type-dependent prognostic associations, yet in solid tumors is generally associated with better overall survival^46^. Altered gal-9 expression is not limited to neoplastic cells; gal-9 has been shown to be upregulated on macrophages and exhausted lymphocytes to suppress antitumor immunity^47, 48, 49^. In DCs however, the literature is scarce. One study described gal-9 protein levels to be enriched in mature myeloid DCs in colorectal cancer tumors and correlated this finding with favorable histological features, underscoring that DC-derived gal-9 is functionally relevant in human cancer^50^. To the best of our knowledge, our study is the first one to describe the selective loss in CD14^+^cDC2s as well as the mechanisms underlying its altered expression.

Mechanistic studies indicate that deregulated galectin (-1, -3 and -9) expression in the TME is typically driven by transcriptional programs (hypoxia and cytokine-inducible pathways via HIF-1α, NF-κB or STAT), epigenetic alterations and microRNA networks^51^. In our system, however, gal-9 modulation in CD14^+^cDC2s is confined to the cell surface, rapidly reversible after removal of tumor-derived soluble factors (within 18 h), and transcription-independent, indicating that the altered gal-9 binding at the DC surface is uncoupled from gal-9 transcript levels. Our observations align with studies showing that transient removal of extracellular galectins with lactose acutely diminishes surface gal-9 without altering its expression, as shown by intact intracellular pools and mRNA levels^27^. Concretely, formation of galectin lattices and control of galectin extracellular functions is heavily dependent on glycome remodeling and altered glycan density and pattern^24^. Notably, we observed no reduction in gal-3 surface levels upon CM exposure, which further supports the existence of gal-9-targeted mechanisms driven by specific tumor-induced glycome changes, consistent with altered multivalency, type of glycan where this glycoepitope is exposed, or ability to crosslink receptors into a lattice for each galectin.

The tumor secretome comprises a vesicular fraction (including exosomes, macrovesicles, apoptotic bodies and oncosomes) and a soluble fraction (metabolites, lipids, nucleic acids, and proteins), all of which have been implicated in tumor-mediated immune evasion^52^. To identify which component drives gal-9 loss, we fractionated tumor-CM by molecular weight and found that only the >100 kDa fraction, comprising glycoconjugates, such as proteoglycans, mucins, or glycoproteins, EVs or proteins complexes reduced gal-9 surface levels. Although matrix metalloproteases (MMPs) have been shown to cleave gal-9 in the TME, the restriction of gal-9 decreased binding to the CD14^+^cDC2 subset argues against an unselective proteolytic mechanism, such as melanoma-derived MMP, as such mechanism should impact all DC subsets^53^. Therefore, we hypothesize that tumor factors could remodel the glycan structures to which gal-9 binds on CD14⁺ cDC2s, making the DC glycome less favorable for gal-9 interactions. This is supported by the fact that altered glycosylation is a hallmark immune evasion strategy in cancer, where changes in tumor glycans are sensed by immune cells via lectins and often translated into inhibitory immune programs^18^. Concomitantly, aberrant expression of glycan motifs such as GlcNAc (N-acetylglucosamine), NeuAc (N-acetylneuraminic acid), TF antigen (Thomsen-Friedenreich antigen) and fucose at the surface of melanoma cells induced DC dysfunction via lectin recognition that could be reversed by interfering with the glycan/lectin axis. These altered glycan patterns have also been associated with poor disease outcome in melanoma patients, underscoring the biological implications of the tumor-associated glycome changes in tumor-residing DCs^54^. Additionally, cDC1s, cDC2s and pDCs may exhibit distinct recognition of melanoma-associated glycan motifs (Gal, Man, GalNAc/Tn, sTn, Fuc, GlcNAc) likely mediated by differential expression of glycan-binding proteins, suggesting that tumor cells might exert glycan-dependent subset-specific (dys)regulation^55^. However, alterations in the DC-glycome during melanoma progression and how it shapes galectin binding and function remains essentially unexplored. Here we found that sialic acids, GalNAc and fucosylated motifs at the surface of cDC2 were reduced after tumor exposure, and that GalNAc decrease mirrored gal-9 loss induced by the >100kDa fraction. This suggests that remodeling of GalNAc-containing glycans could weaken multivalent gal-9 interactions or redirect gal-9 to alternative ligands, providing a mechanistic link between tumor-driven DC glyco-reprogramming and selective gal-9 decreased binding. Future glycoproteomic studies could identify glycan changes within specific gal-9 ligands at the DC surface to better map gal-9-dependent membrane receptor binding upon exposure to the tumor secretome.

Several non-mutually exclusive mechanisms could explain how melanoma-derived high-molecular-weight factors reduce GalNAc-containing epitopes on DCs. One possibility is active enzymatic remodeling of surface GalNAc structures through decreased glycosyltransferases involved in biosynthesis of GalNAc-containing motifs, or increased glycosidase activity within the TME^56^. For instance, α-N-acetylgalactosaminidase (NAGA) is upregulated in serum of melanoma patients^57^, and this enzyme could be present in melanoma vesicles. Given that decrease of GalNAc motifs is observed in the high MW secretome fraction (>100kDa), and considering that NAGA has a MW of 46-47 kDa and its mRNAs ∼7 kDa, our data support the presence of this enzyme as cargo in extracellular vesicles (EVs) or larger multimolecular complexes, which has been previously described for other glycan-remodeling enzymes in melanoma as a delivery mechanism^58, 59, 60^. Another plausible mechanism could be melanoma-induced transcriptional reprogramming of DCs that alters the expression of enzymes involved in glycan biosynthesis and degradation. Tumor-derived EVs and soluble factors have been shown to modulate gene expression programs in myeloid cells^61, 62^, and could therefore influence the glycosylation machinery in DCs, including glycosyltransferases responsible for the synthesis or modification of GalNAc-containing structures (e.g. members of the GalNAc-transferase family or other enzymes such as those related to ABO-like glycosylation pathways). Such changes could shift the balance of glycan biosynthesis toward reduced presentation of terminal GalNAc epitopes at the DC surface. Finally, dysregulation of cell surface glycoproteins carrying these GalNAc motifs remains a possible alternative hypothesis. Altogether, these observations support a model in which melanoma-derived EVs or high-molecular-weight multimolecular complexes deliver glycan-remodeling enzymes and/or regulatory RNAs to intratumoral DCs, culminating in active loss of GalNAc-containing epitopes (reduced DBA reactivity) and diminished gal-9 binding.

Despite reduced gal-9 binding upon tumor exposure, r-gal-9 supplementation of tumor-conditioned CD14⁺cDC2 retained sufficient binding to partially restore immunostimulatory function, reflected by reduced Treg induction in allogeneic T cell co-cultures to near non-tumor-exposed levels. This finding is consistent with work in food allergy showing that exogenous gal-9 can recalibrate dysfunctional human DCs and restore their tolerogenic Treg-inducing capacity in patients with food allergy^63^. In contrast, in tumor settings our finding contradicts previous T cell studies which describe that gal-9 promotes the generation, stability and suppressive function of Foxp3⁺ Tregs^64, 65, 66^. Notably, these reports mostly used gal-9 exogenously treated T cells or studied gal-9 produced by Tregs themselves in the context of strong TGF-β and Tim-3/CD44 signaling. Additionally, in vivo studies employing anti-gal-9 with PD-1/PD-L1 blockade did not report a Treg depletion or functional impairment^67^. On the contrary, here we specifically modulated gal-9 on melanoma-conditioned DCs. Thus, overall, this supports a model in which gal-9 fine-tunes the DC-T cell axis in a context and cell-dependent manner rather than uniformly enhancing Treg responses.

In conclusion, our data collectively highlighted a previously uncovered gal-9 loss in human CD14^+^cDC2 cells accompanied by acquisition of immunosuppression phenotype. This loss was mirrored by melanoma-induced glycan changes on the DC membranes, which correlated with altered r-gal-9 binding. Notably, by rescuing tumor-induce gal-9 loss, we could modulate Treg abundance, suggesting that targeting gal-9 may have important implications for tumor escape via Treg-dependent mechanisms.

