## Supplementary Figures for "Melanoma suppresses galectin-9-glycan axis in dendritic cells and galectin-9 restoration limits T regulatory cell expansion"

**Supplementary Figure 1**

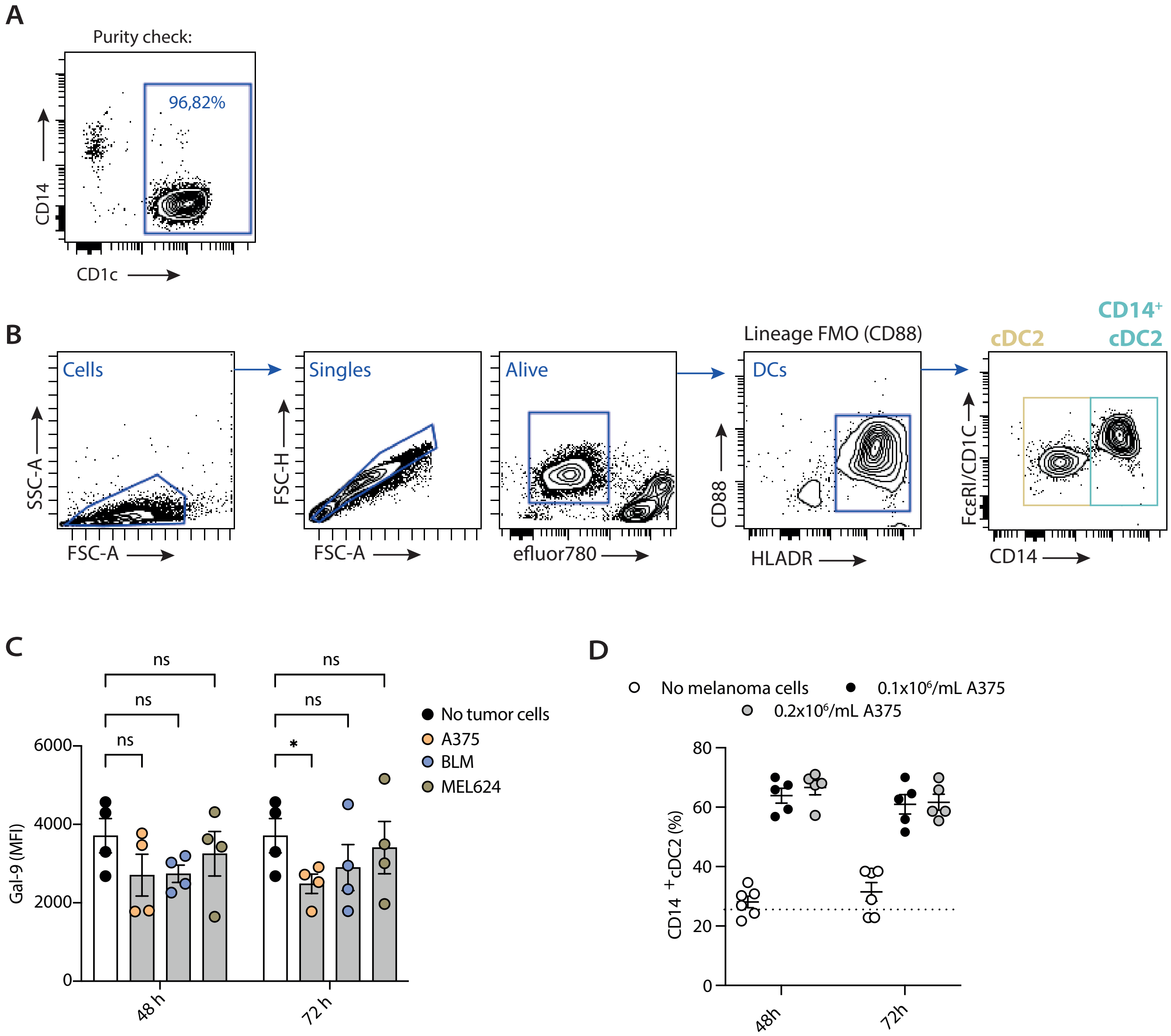

Supplementary Figure 2

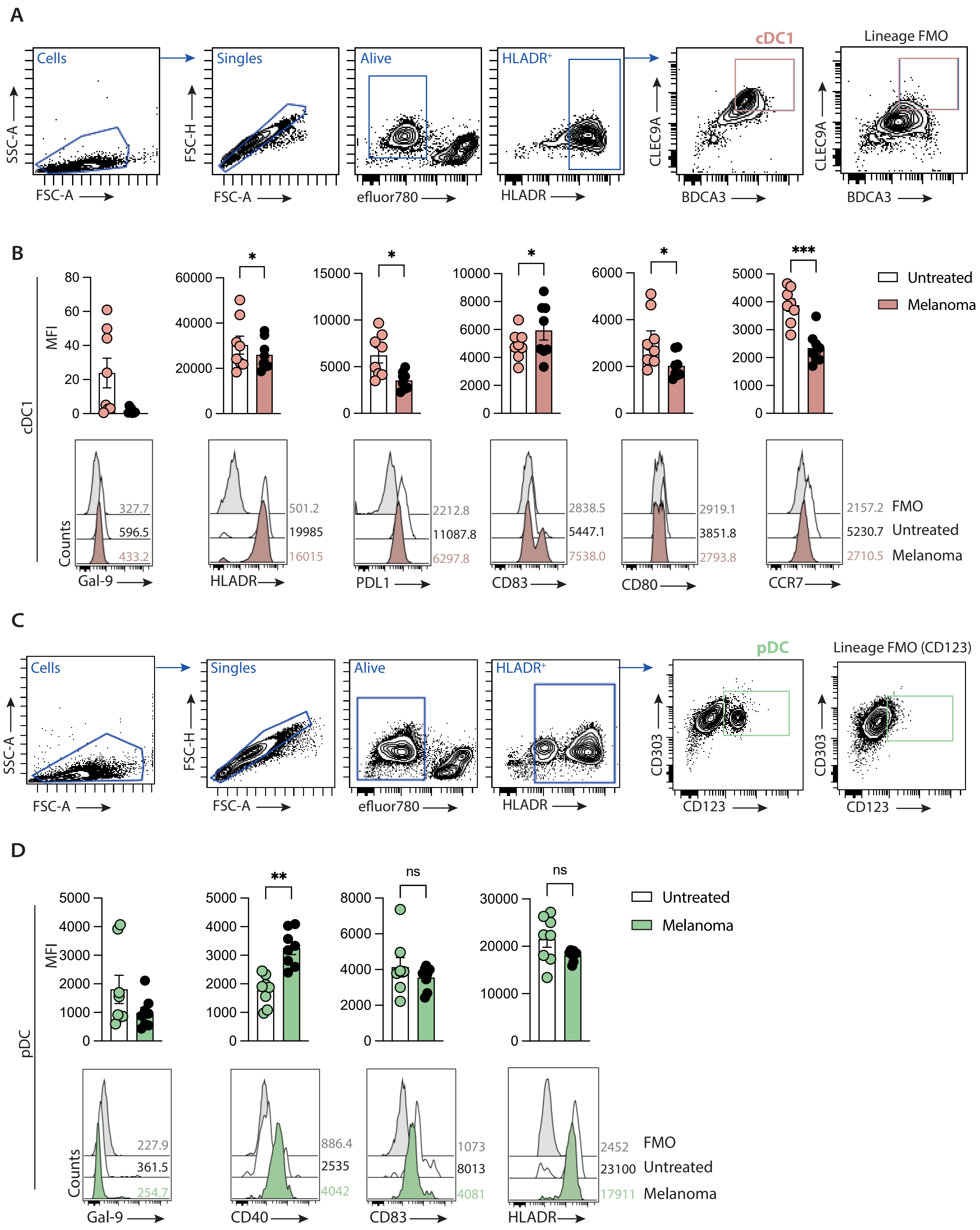

### Supplementary Figure 3

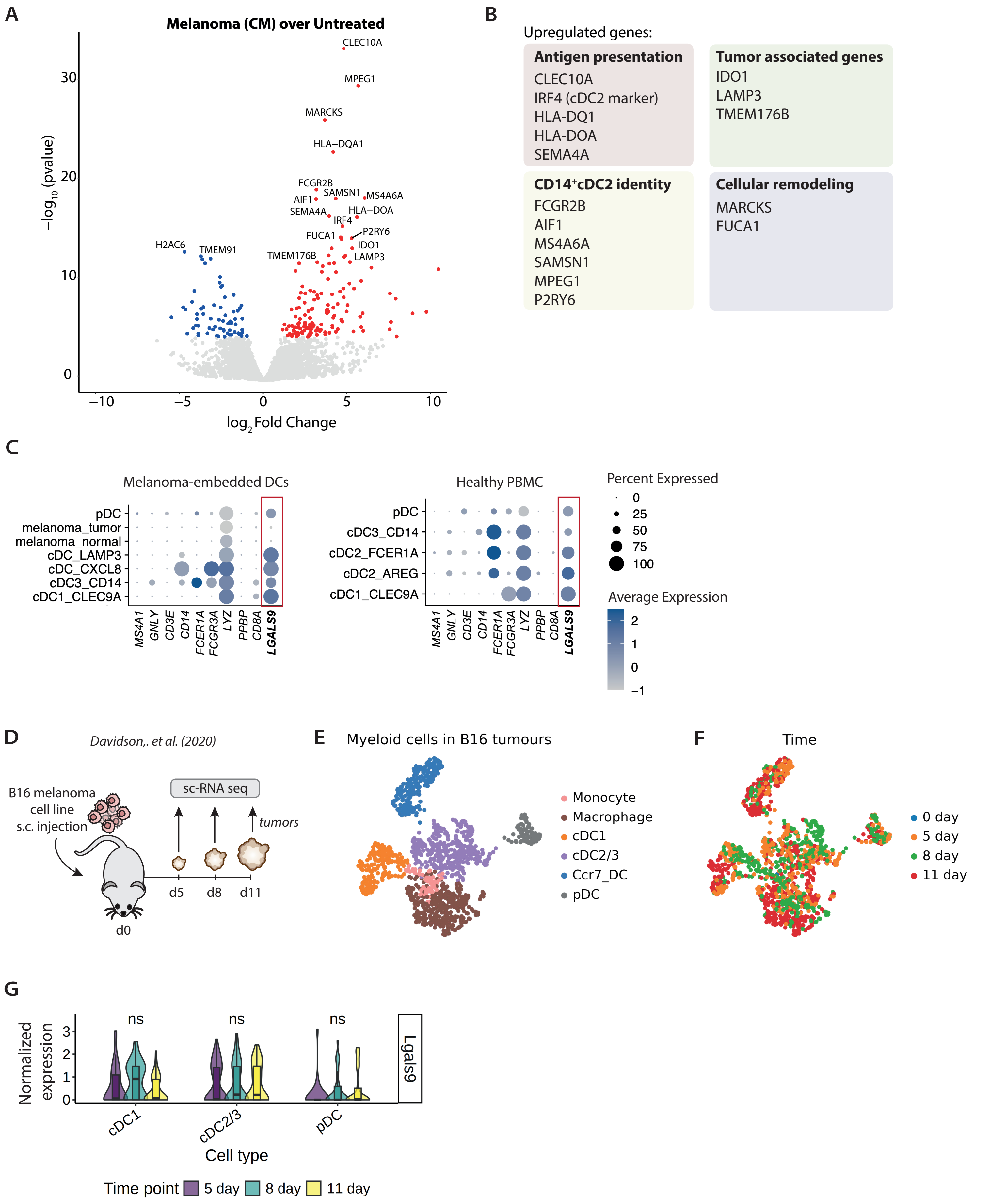

**Supplementary Figure 4**

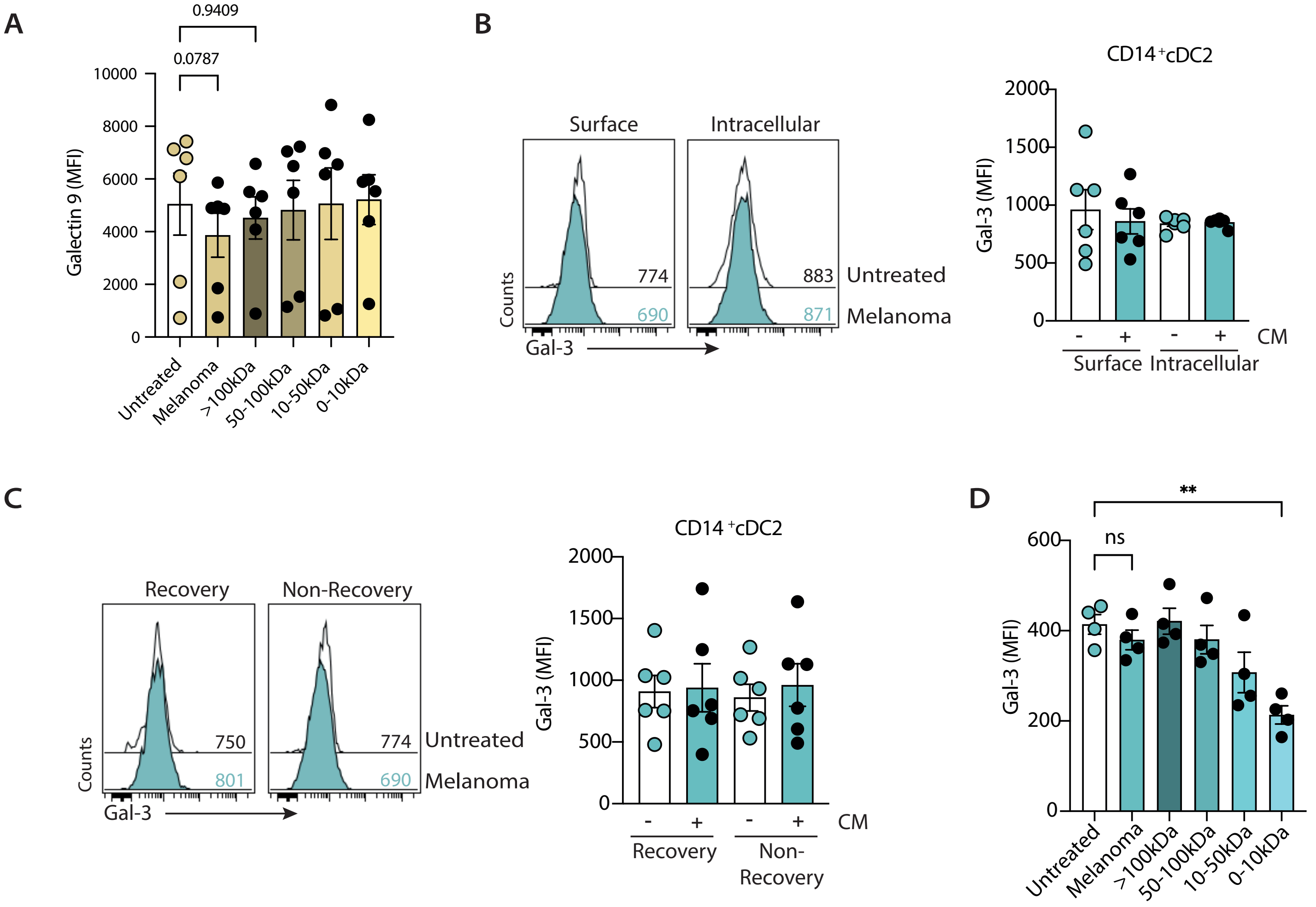

**Supplementary Figure 5**

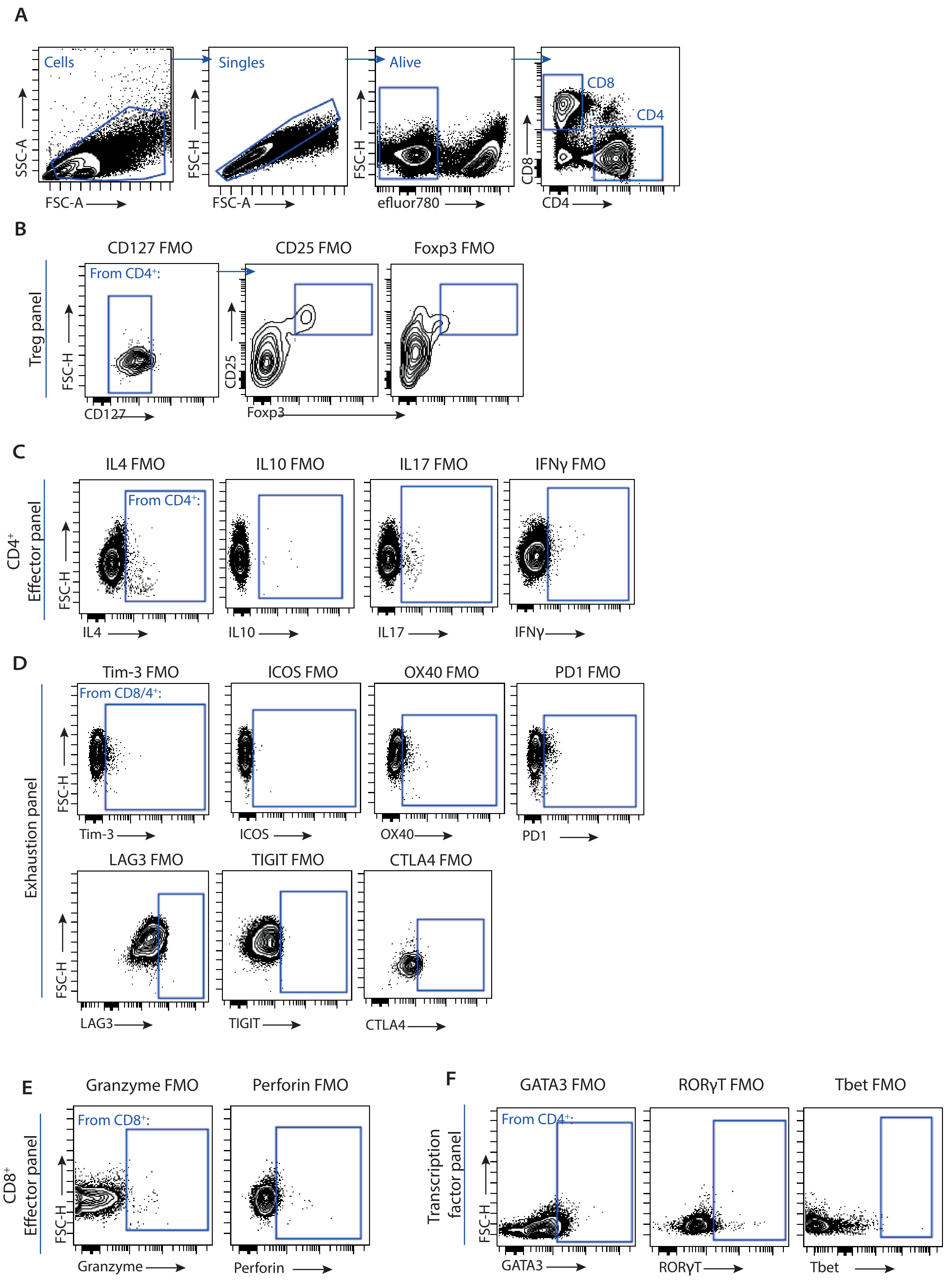

**Supplementary Figure 6**

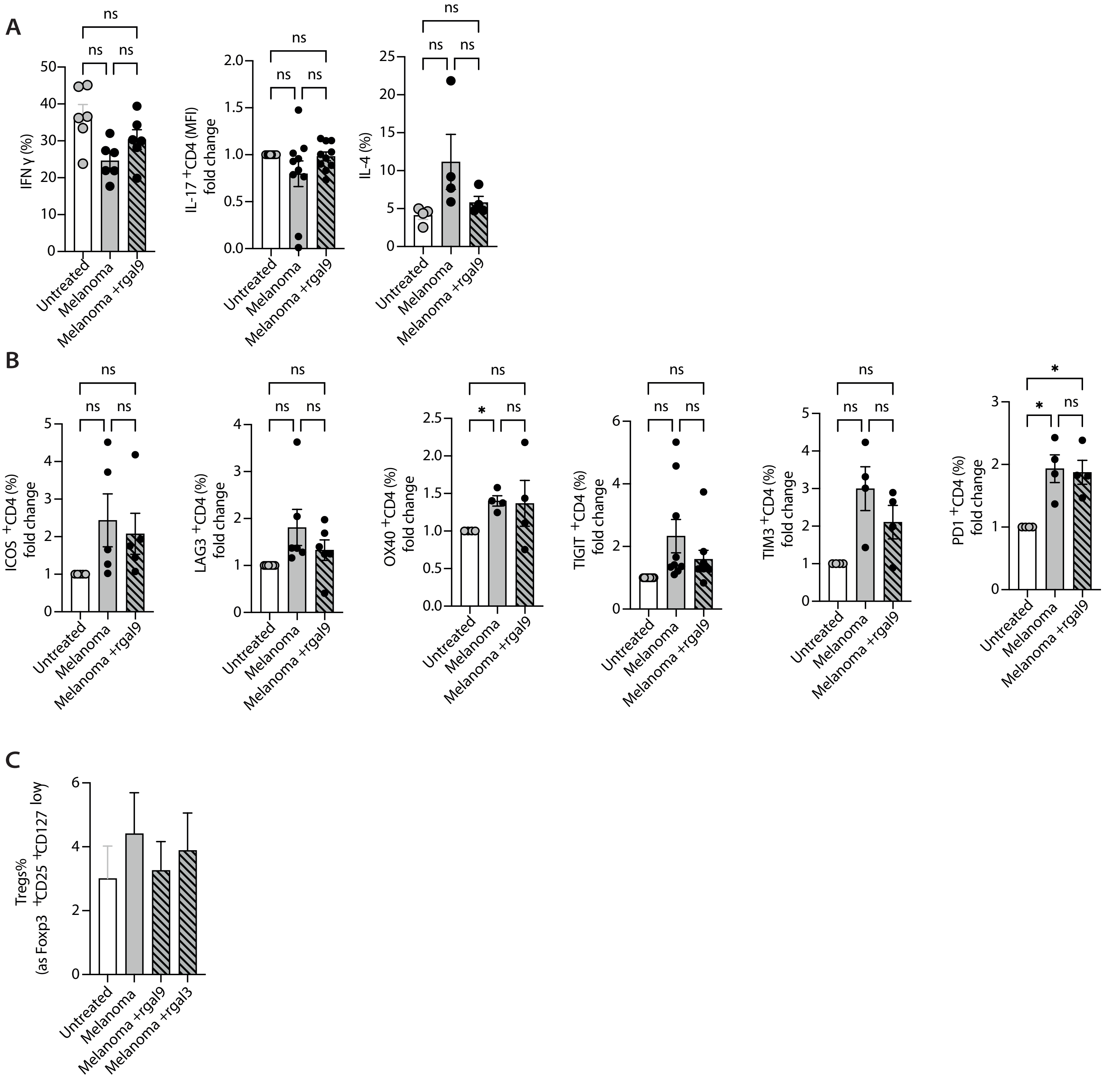

**Supplementary Figure 7**

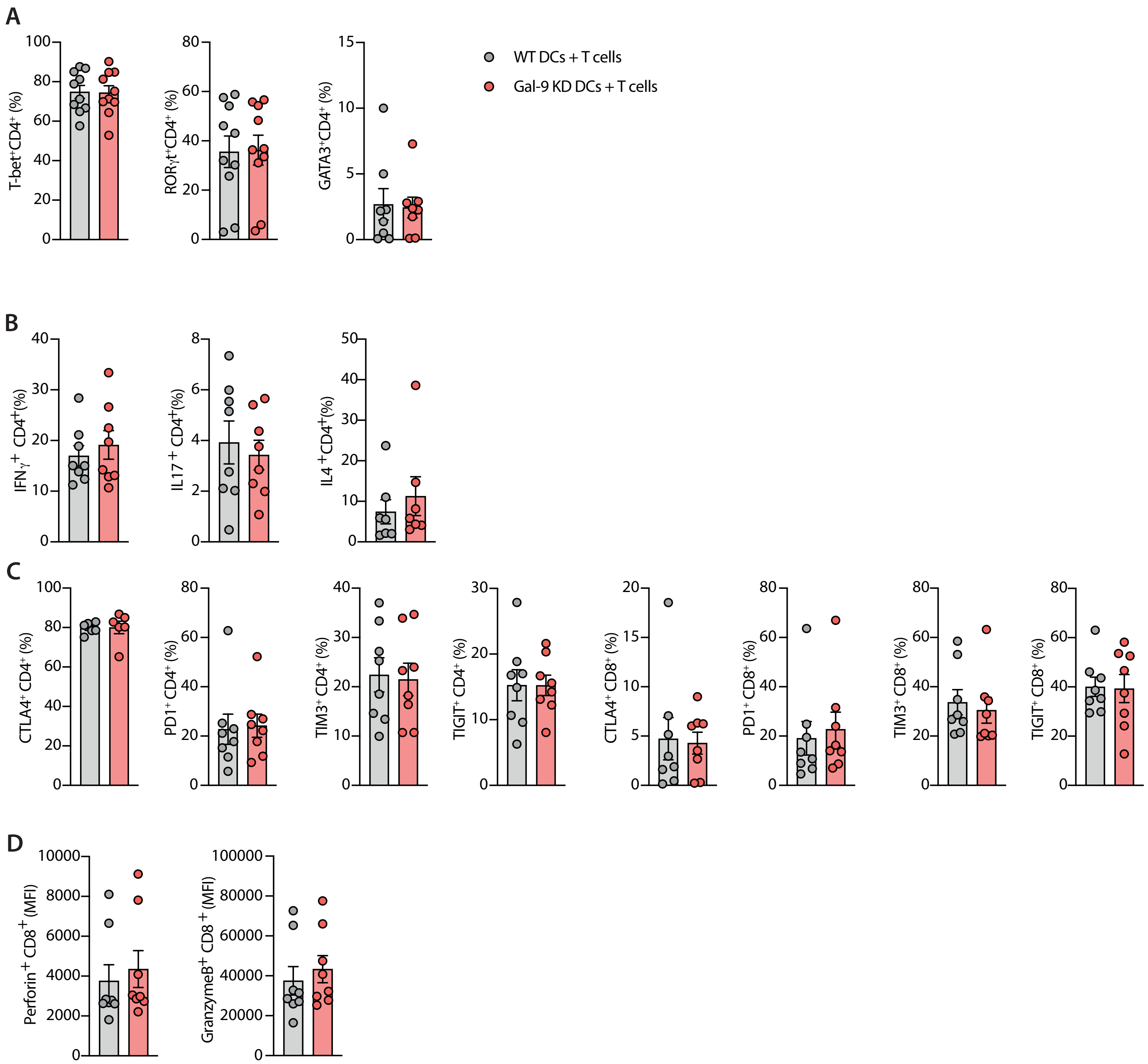

### Supplementary figure legends

**Supplementary Figure 1. Melanoma-derived CM decreases surface gal-9 in cDC2s. A.** Purity check of sorted cDC2 population based on CD1c and CD14 expression, purity indicated in the top right part of the graph. **B.** Sequential gating for cDC2/CD14$^+$cDC2. Lineage FMO controls shown for CD88. Gates used are displayed by a square. **C.** Gal-9 surface expression in cDC2s upon incubation for 48 or 72 hours with different melanoma cell line-derived CM (A375, BLM and MEL624). **D.** Quantification (%) of CD14$^+$cDC2 cells obtained after 48 or 72 hours of normal media ("no melanoma cells", white) or melanoma-CM treatment (at different A375 seeding concentrations, 0,1-0,2x10$^6$ cells/mL, black and gray). Data shown as mean ± SEM. Each dot represents an independent donor. Statistical significance displayed as ns $p > 0.05$ and *$p < 0.05$.

**Supplementary Figure 2. Gating strategy for DC subset identification and phenotypic characterization of DC subsets treated or not with melanoma-CM**. Sequential gating for cDC1 (**A**) and pDC (**C**) subset identification. Lineage FMO controls shown for CLEC9A and CD123. Gates used are displayed by a square. Representative flow cytometry histogram (down) and MFI quantification (up) (n=8) for expression of gal-9, HLA-DR, PD-L1, CD83, CD80, and CCR7 on cDC1 subset (**B**) and gal-9, CD40, CD80 and HLADR on pDCs (**D**). Unstained control or FMO samples are shown in gray, untreated samples are displayed in white and melanoma-CM treated samples in red (for cDC1) and green (for pDCs). Numbers in histograms indicate the MFI value of each histogram. Data shown as mean ± SEM. Each dot represents an independent donor. Statistical significance assessed by paired student T test. ns $p > 0.05$, *$p < 0.05$, **$p < 0.01$, ***, $p < 0.001$.

**Supplementary Figure 3. Gal-9 changes are independent of transcriptional regulation (in murine and human data). A**. Volcano plot showing differentially expressed genes in human CD14$^+$ cDC2s treated with melanoma-CM relative to untreated controls. Up-regulated genes are shown in red, down-regulated in blue and non-significant genes in gray. **B.** Core differentially expressed genes in CD14$^+$ cDC2s after melanoma treatment are highlighted and categorized into different gene groups, summarizing the functional programs where they are involved (antigen presentation, tumor-associated genes, genes linked to CD14$^+$ cDC2 identity and cellular remodelling). **C.** Dot plot showing gene expression patterns across DC subsets in melanoma-exposed (left) and healthy PBMCs (right). Each dot represents the expression of a given gene (columns) in a specific cell type. Dot size indicates the percentage of cells expressing the gene, while dot color represents the scaled average expression level (blue = higher expression, gray = lower expression). Gal-9 (*LGALS9*) gene expression is framed in red. **D.** Schematic overview adapted from Davidson et al. (2020) showing the experimental design of the B16 melanoma mouse model analyzed by sc-RNA-seq [72]. Mice were subcutaneously (s.c.) injected with B16 melanoma cells at day 0, and tumors -and adjacent lymph nodes- were subsequently harvested on days 5, 8 and 11 post-injections. Sc-RNA-seq was performed on these tissues to identify *lgals9* transcriptional changes in DCs within the TME during melanoma progression. **E.** UMAP projection of myeloid cells isolated from B16 tumors, colored by annotated cell populations, including monocytes, macrophages, cDC1, cDC2/3, Ccr7$^+$ dendritic cells (Ccr7_DC), and pDC. **F.** The same UMAP colored by time. **G.** Violin plots showing normalized expression of *lgals9* across dendritic cell subsets (cDC1, cDC2/3, and pDC) at days 5 (purple), 8 (green), and 11 (yellow) following B16 melanoma implantation.

**Supplementary Figure 4. Melanoma-CM components do not modulate galectin-9 or -3 expression in CD14$^+$cDC2 or CD14$^-$cDC2. A.** Quantification of gal-9 MFI (n=6) of untreated (white) or fractionated CM-exposed CD14$^-$cDC2s (gradients of yellow). **B.** Representative flow cytometry histograms of surface (left) and intracellular (right) gal-3 expression in untreated (white) or melanoma-exposed CD14$^+$cDC2 cells (blue). Quantification of gal-3 MFI (n=6) for surface and intracellular staining shown on the right. **C.** Representative histograms of gal-3 expression after recovery (left) or non-recovery (right) conditions in untreated (white) or melanoma-exposed CD14$^+$cDC2 cells (blue). Quantification of gal-3 MFI (n=6) shown on the right. **D.** Quantification of gal-3 MFI (n = 6) in CD14$^+$cDC2 cells exposed to melanoma-CM or CM fractions (>100 kDa, 100-50 kDa, 50-10 kDa, 10-0 kDa). Numbers in histograms indicate the MFI value of each histogram. Data shown as mean ± SEM. Each dot represents an independent donor. Statistical significance assessed by two-way ANOVA with Šídák's multiple comparisons: ns $p > 0.05$, *$p < 0.05$, **$p < 0.005$, **** $p < 0.00005$.

**Supplementary Figure 5. Gating strategy for T cell phenotyping. A.** Sequential gating for CD4$^+$ and CD8$^+$ T cell identification. **B.** Gating for Tregs based on CD127, CD25, and Foxp3 expression (FMO shown). **C.** Gating for cytokine-producing CD4$^+$ T cells using FMO controls for IL-4, IL-10, IL-17, and IFNγ. **D.** Gating for activation and exhaustion markers including Tim-3, ICOS, OX40, PD-1, LAG3, TIGIT, and CTLA-4 using FMO controls. **E.** Gating for cytotoxic markers in CD8$^+$ T cells: Granzyme B and Perforin using FMO controls. **F.** Gating for transcription factors: STAT3, RORγT, and Tbet using FMO controls. Gates used are displayed by a square.

**Supplementary Figure 6. Effect of melanoma-CM-treated DCs on T cell phenotype and function. A.** IFNγ$^+$, IL-17A$^+$ and IL-4A$^+$ CD4$^+$ T cell frequency across conditions (n=4-10). **B.** Expression of activation and exhaustion markers in CD4$^+$ T cells across conditions (n=4-8). IL-17 and exhaustion markers are shown as the relative frequencies of melanoma-treated cells compared to untreated. **C.** Quantification of Treg frequency (Foxp3$^+$CD25$^+$CD127$^{low}$) under untreated, melanoma-conditioned, and melanoma-conditioned plus r-gal-9 or r-gal-3 conditions (n=7). Untreated conditions are depicted in white, melanoma-conditioned in gray, and melanoma-conditioned plus r-gal-9 in stripped gray. Data shown as mean ± SEM; each dot represents an independent donor. Statistical significance assessed by two-way ANOVA with Šídák's multiple comparisons: ns $p > 0.05$, *$p < 0.05$.

**Supplementary Figure 7. Impact of gal-9 KD in DCs on T cell phenotype and function. A.** Frequency (%) of T cell helper subset transcription factors (T-bet, RORγt and GATA3) in CD4$^+$ T cells co-cultured with WT DCs (gray) or gal-9 KD DCs (red) (n=10). **B.** Cytokine-producing CD4$^+$ T cells: IL-4$^+$, IFNγ$^+$, and IL-17A$^+$ frequencies across conditions (n=8). **C.** Expression of exhaustion markers in CD4$^+$ and CD8$^+$ T cells co-cultured with WT DCs or gal-9 KD DCs (n=8). **D.** Cytotoxic granule marker expression (Perforin and Granzyme B) in CD8$^+$ T cells under the same conditions (n=8). Data shown as mean ± SEM; each dot represents an independent donor. Statistical significance assessed by paired student T test.
